# Genome-wide gene expression profiling reveals that cuticle alterations and P450 detoxification are associated with pyrethroid resistance in *Anopheles arabiensis* populations from Ethiopia

**DOI:** 10.1101/451336

**Authors:** Eba Alemayehu Simma, Wannes Dermauw, Vasileia Balabanidou, Simon Snoeck, Astrid Bryon, Richard M. Clark, Delenasaw Yewhalaw, John Vontas, Luc Duchateau, Thomas Van Leeuwen

## Abstract

**BACKGROUND:** Vector control is the main intervention in malaria control and elimination strategies. However, the development of insecticide resistance is one of the major challenges for controlling malaria vectors. *Anopheles arabiensis* populations in Ethiopia showed resistance against both DDT and the pyrethroid deltamethrin. Although a L1014F target-site resistance mutation was present in the voltage gated sodium channel of investigated populations, the levels of resistance and biochemical studies indicated the presence of additional resistance mechanisms. In this study, we used genome-wide transcriptome profiling by RNAseq to assess differentially expressed genes between three deltamethrin and DDT resistant *An. arabiensis* field populations (Tolay, Asendabo, Chewaka) and two susceptible strains (Sekoru and Mozambique).

**RESULTS:** Both RNAseq analysis and RT-qPCR showed that a glutathione-S-transferase, *gstd3*, and a cytochrome P450 monooxygenase, *cyp6p4*, were significantly overexpressed in the group of resistant populations compared to the susceptible strains, suggesting that the enzymes they encode play a key role in metabolic resistance against deltamethrin or DDT. Furthermore, a gene ontology enrichment analysis showed that expression changes of cuticle related genes were strongly associated with insecticide resistance, although this did not translate in increased thickness of the procuticle.

**CONCLUSION:** Our transcriptome sequencing of deltamethrin/DDT resistant *An. arabiensis* populations from Ethiopia suggests non-target site resistance mechanisms and pave the way for further investigation of the role of cuticle composition in resistance.

## 1. Introduction

Vector control is the main intervention in malaria control and elimination strategies. Indoor residual spraying and insecticide-treated nets have made substantial contributions to the reduction of malaria incidence. ^1, 2^ However, the development of insecticide resistance in the major anopheline malaria vectors threatens the global effort to control malaria.^3–5^ Some *Anopheles gambiae* mosquito populations now show resistance to all insecticide classes and the strengths, and the impact of resistance is escalating every year.^6^ High levels of insecticide resistance have also been reported for *An. arabiensis* in many countries, including Ethiopia ^7–11^, where we recently surveyed several populations across the country and showed that these *An. arabiensis* populations exhibited countrywide resistance against DDT and the pyrethroid deltamethrin.^12^

Understanding the molecular mechanisms underlying resistance has the potential to aid developing of strategies to prevent and/or delay the spread of insecticide resistance in malaria vectors including *An. arabiensis*.^13^ Resistance mechanisms can be classified into two mechanisms. Alterations of the target-site, for example by point mutations, which reduce the susceptibility to pesticides, are known as toxicodynamic mechanisms. Increased detoxification, decreased penetration, sequestration or increased excretion of insecticides through qualitative or quantitative changes of enzymes/proteins are known as toxicokinetic mechanisms. ^14, 15^ Finally, behavioral mechanisms, such as avoidance of insecticide exposure, have been proposed as a third resistance mechanism, but up until now no conclusive evidence has been reported that supports this type of resistance mechanism. ^16^

Several variants in the *knockdown resistance* (*kdr*) gene, encoding the voltage-gated sodium channel (VGSC), have been shown to be, or are associated with, pesticide resistance in malaria vectors. The VGSC is the target-site of pyrethroids and DDT. Pyrethroids are currently the main insecticide class used to control malaria vectors.^3^ Several mutations in the VGSC that confer pyrethroid resistance have been reported in *Anopheles* vectors. ^17^ These result in the substitution of leucine 1014 (TTA) to phenylalanine (TTT) (*kdr* L1014F) or to serine (TCA) (*kdr* L1014S). Additionally, a N1575Y mutation in VGSC was reported to have a synergistic effect with the L1014F mutation, and has so far has been observed in *An. gambiae* and *An. coluzzi* species ^18, 19^, but not in *An. arabiensis*. A G119S mutation in the target-site of organophosphates and carbamates, acetyl-choline esterase 1 (AChE1), has been described for resistant *An. gambiae* populations in West-Africa ^20, 21^, as well as more recently in *An. arabiensis* ^22^. Last, the GABA-gated chloride channel is known as the target of cyclodienes and resistance mutations (A301S, V332I and T350S) in the gene encoding this channel (*resistance to dieldrin, rdl*) were shown to confer cyclodiene resistance in anopheline populations. ^23, 24^

In addition to target-site resistance, cytochrome P450 monooxygenases (P450s) are the most important enzyme family involved in toxicokinetic resistance mechanisms of insects to pyrethroids. ^3, 17^ In many resistant strains of *Anopheles* species, P450s have been shown to be overexpressed and able to metabolize pyrethroids^25–28^. For example, the P450s encoded by *cyp6m2* and *cyp6p3*, the most widely over-expressed P450s in pyrethroid resistant field populations of *An. gambiae*, are both capable of metabolizing pyrethroids.^25, 26, 28^ In some cases, overexpressed P450s also confer resistance to insecticide classes other than pyrethroids. For example, *An. gambiae* CYP6M2 and CYP6P3 can metabolize the organochlorine DDT and the carbamate bendiocarb, respectively.^29, 30^ Further, the P450s CYP6P4 and CYP4G16 have been associated with pyrethroid resistance in *An. arabiensis*.^31, 32^ Ibrahim *et al.* 2016 showed that CYP6P4 plays a key role in pyrethroid resistance of *An. arabiensis* populations from Central Africa (Chad), and that it can metabolize the pyrethroids permethrin, bifenthrin and λ-cyhalothrin, but not deltamethrin.^31^ Another P450, CYP4G16, has been associated not with pyrethroid metabolism directly, but with the increased biosynthesis of epicuticular hydrocarbons that delay insecticide uptake.^33^ Notably, in addition to epicuticular hydrocarbon enrichment, compositional changes of the cuticle have also been associated with insecticide resistance and several genes have been associated with the phenomenon.^34^ Finally, glutathione-S-transferases (GSTs) are also important enzymes involved in toxicokinetic resistance mechanisms against pyrethroids.^35–38^ For example, Riveron *et al.* 2014, 2017, showed that allelic variation and higher transcription of GSTe2 confers resistance against permethrin in an *An. gambiae* population of Benin.^35, 36^ However, the role of such GST-based quantitative and qualitative changes has not yet been investigated in pyrethroid resistant *An. arabiensis*.

Recently, we surveyed several *An. arabiensis* populations from Ethiopia and showed that all these populations exhibited resistance against DDT and the pyrethroid deltamethrin.^12^ The frequency of the target-site resistance mutation L1014F in the VGSC was high in some populations, but resistance levels suggested additional resistance mechanisms, as has been observed in other *Anopheles* species.^39–41^ In contrast to *An. gambiae*, only few genome-wide gene expression studies have investigated insecticide resistance in *An. arabiensis* populations^32, 42, 43^, and none of them focused on Ethiopian populations. In this study, we performed genome-wide transcriptome profiling (RNAseq, Illumina platform) with three resistant populations and two reference susceptible *An. arabiensis* strains from Ethiopia and identified candidate genes and mechanisms for insecticide resistance in this important malaria vector.

## 2. Materials and methods

### 2.1. Mosquito populations

*An. arabiensis* larvae were collected in the South-West part of Ethiopia from a range of breeding sites: Asendabo (ASN), Chewaka (CHW) and Tolay (TOL) (Figure S1). Larvae were reared to adults on site in rooms with standard conditions of 25 ± 2°C and a relative humidity of 80 ± 10% for all three respective sites. Larvae were fed with dog biscuits and brewery yeast whereas adults were provided a 10% sucrose solution soaked into cotton pads^44^. ASN, CHW and TOL were previously shown to be resistant against deltamethrin and DDT^12^. Two laboratory strains served as pesticide susceptible populations: an Ethiopian strain (Sekoru (SEK)), previously described in Alemayehu *et al.* 2017, and a strain from Mozambique (MOZ) previously described in Witzig *et al.* 2014.^12, 45^ Both laboratory strains were reared in a similar way as the three Ethiopian populations collected from the field.

### 2.2. RNA extraction

Batches of five 3-5-day-old, non-blood-fed *An. arabiensis* female mosquitoes from each population (ASN, CHW or TOL) or strain (SEK, MOZ) were preserved in RNAlater (Ambion, Thermo Fischer Scientific) in a 1.5ml Eppendorf tubes. In total, between eighty to hundred adult females were collected for each population/strain. All tubes were stored at −80°C. The field-collected samples were transported on dry ice to the laboratory of Agrozoology, Department of Plants and Crops (University of Ghent, Belgium). Total RNA was extracted from batches of ten female mosquitoes using the RNAqueous®-4PCR Total RNA isolation Kit (Ambion, Thermo Fischer Scientific). RNA was treated with DNase1 and DNase was inactivated according to the instructions for the RNAqueous®-4PCR Kit. Four biological replicates were included for each population or laboratory strain. Total RNA samples were quantified with a DeNovix DS-11 spectrophotometer (DeNovix, USA) and visualized by running an aliquot on a 1% agarose gel.

### 2.3. RNAseq library preparation and sequencing

Illumina libraries were constructed with the TruSeq Stranded mRNA Library Preparation Kit with polyA selection (Illumina, USA), and the resulting libraries were sequenced on an Illumina HiSeq 2500 instrument to generate strand-specific, paired-end reads of length 125 bp (HiSeq SBS Kit v4 sequencing reagents). Library construction and sequencing was performed at the High-Throughput Genomics and Bioinformatic Analysis Shared Resource at Huntsman Cancer Institute (University of Utah, Salt Lake City, UT, USA). According to FastQC version 0.11.4^46^ no reads were tagged as poor quality. The RNAseq expression data generated during the current study are available in the Gene-Expression Omnibus (GEO) repository with accession number GSE121006 (reviewer token:qjkreseaxnwjluf).

### 2.4. Differential expression and Gene Ontology (GO) enrichment analysis

All reads were aligned to the nuclear genome^47^ and mitochondrial genome (GenBank accession: NC_028212) of *An. arabiensis* using HISAT2^48^ and the following options “‐‐maxintronlen 75000 ‐‐rna-strandness RF ‐‐known-splicesite-infile splicesites.txt”. The “splicesites.txt” file was generated from the gene transfer format (GTF) files of the nuclear and mitochondrial genome of *An. arabiensis* using a script accompanying the HISAT2 software (hisat2_extract_splice_sites.py). For the nuclear genome, the AaraD1.6 GTF annotation file was used (13830 genes of which 13452 are protein coding genes, released 25 April 2017 at VectorBase^49^, https://www.vectorbase.org/organisms/anophelesarabiensis/dongola/aarad16); for the mitochondrial genome a GTF file was generated from the GenBank file (NC_028212.1) using the bp_genbank2gff3.pl and gffread script included in the BioPerl (http://bioperl.org) and Cufflinks package^50^, respectively (see File S1 for the *An. arabiensis* GTF used for mapping). Resulting BAM files were subsequently sorted by read name using SAMtools version 1.5.^51^ Next, read counts per gene were obtained using the htseq-count script included in the HTSeq package, version 0.9.0^52^, with the following settings “-i gene_id ‐t exon ‐f bam ‐s reverse.”. Differential gene expression (DE) analyses were performed using DESeq2 (version 1.12.2).^53^ Differentially expressed genes (DEGs), as assessed with a fold change (FC) ≥ 2 and Benjamini-Hochberg adjusted *p*-value (FDR) < 0.05, were determined between each resistant population and each susceptible strain (six comparisons in total: ASN vs. MOZ, ASN vs. SEK, CHW vs. MOZ, CHW vs. SEK, TOL vs. MOZ and TOL vs. SEK, Figure 2). For the DESeq2 output of all comparisons, a GO enrichment analysis was performed using the Bioconductor package GOSeq (version 1.24.0) with FDR = 0.05. The GOSeq package takes into account gene selection bias due to differences in gene (median transcript) length. GO terms for *An. arabiensis* nuclear genes were downloaded from VectorBase (https://www.vectorbase.org)^49^ using BioMart, while the GO terms for *An. arabiensis* mitochondrial genes were identified using InterProScan version version 5.25-64.0, available at the EMBL-EBI website (https://www.ebi.ac.uk/interpro/interproscan.html).

### 2.5. Principal component analysis and gene expression heatmap

A Principal Component Analysis (PCA) was performed as described by Love *et al.* 2015.^54^ Briefly, read counts were first normalized using the regularized-logarithm (rlog) transformation implemented in the DESeq2 (version 1.12.2) R-package. A PCA was then performed using the stats (version 3.3.0), ggbiplot (version 0.55) and ggplot2 (version 2.2.0) R-packages with the 1000 most variable genes across all RNAseq samples and the ggbiplot argument ellipse.prob set to 0.95. Gene expression patterns of cuticle related genes were visualized with heatmaps generated with the relative transcript levels (fold changes) of four DE analyses (ASN vs. SEK, CHW vs. SEK, TOL vs. SEK and MOZ vs. SEK) with the limma (version 3.28.21) and gplots (version 3.0.1) packages in the R environment. Cuticle related genes were selected based on the following InterPro domains: IPR000618 (Insect cuticle protein), IPR031311 (Chitin-binding type R&R consensus), IPR002557 (Chitin binding domain), IPR004302 (Cellulose/chitin-binding protein, N-terminal), IPR004835 (Chitin synthase), IPR031874 (Adult cuticle protein 1) and IPR22727 (Pupal cuticle protein C1).

### 2.6. RT-qPCR validation

A subset of *An. arabiensis* DEGs was selected for RT-qPCR validation. Gene specific RT-qPCR primers were designed using Primer3 v.4.1.0.^55^ All primer sequences can be found in Table S1. Total RNA was extracted as described above and cDNA was synthesized with the Maxima First Strand cDNA synthesis for RT-qPCR kit (Fermentas Life Sciences, Aalst, Belgium) starting with 2 μg of total RNA as template. Three biological and two technical replicates were included for each population as well as non-template controls to exclude sample contamination. The RT-qPCR analysis was performed on a Mx3005P qPCR thermal cycler (Stratagene, Agilent Technologies, Diegem, Belgium) with Maxima SYBR Green qPCR Master Mix (2x) and ROX solution (Fermentas Life Sciences) according to the manufacturer’s instructions. qPCR run conditions were: 95°C for 10 m followed by 35 cycles of 95 °C for 15 s, 55 °C for 30 s and 72 °C for 30 s. At the end, a melting curve was generated from 65 °C to 95 °C, 1 °C per 2 s to check for the presence of a single amplicon. Fourfold dilution series of pooled cDNA were used to determine the standard curves and amplification efficiencies for every gene-specific primer pair. Relative expression levels and significant gene expression differences (one-sided unpaired t-test) were calculated with qbase+ version 3.0.^56^

### 2.7. Analysis of mutations involved in insecticide resistance

The presence of mutations involved in *Anopheles* sp. resistance against either DDT, pyrethroids, cyclodienes or organophosphates (I114T and L119F in *gste2* (AARA008732)^36, 40^, L1014C/F/S/W and N1575Y in *vgsc* (AARA016386) (*Musca domestica* numbering^17, 19, 57^), A301S, V332I and T350S in *rdl* (AARA016354) (*Drosophila melanogaster* numbering^24, 58^) and G119S in *AChE1* (AARA010659) (*Torpedo californica* numbering^20, 22^)) was investigated by creating a Variant Call Format (VCF) file from the BAM files employed for analyzing differential gene expression (see above). The BAM files were used as input for SAMtools version 1.4.1^51^ with the following settings “mpileup ‐uf ‐‐output-tags “AD,DP”. Subsequently, the SAMtools output was used as input for BCFtools 1.5.1^51^ with the following settings “call ‐vc”. The effect of single nucleotide polymorphism (SNPs) and small indels on coding sequences in genomic regions were predicted using SNPeff v. 4.3t^59^ with a custom-built *An. arabiensis* coding sequence database (AaraD1.6 annotation for the *A. arabiensis* nuclear genome and NC_028212.1 for the mitochondrial genome) available at VectorBase^49^. Mutation frequencies in target-site genes were calculated based on the frequencies of the reference (“REF”) and alternative (“ALT”) alleles in the allelic depth (“AD”) tag in the SAMtools output.

### 2.8. Cuticle measurements with transmission electron microscopy

The cuticle thickness of mosquito legs from the ASN population and the SEK strain was measured by transmission electron microscopy (TEM), as previously described.^33^ Only the procuticle thickness was measured as the epicuticle layer was abraded in more than 95% of sections during the multiple hexane washes. Only individuals with similar wing size were selected and further analyzed by TEM. Ultra-thin gold sections of the femur leg segment were taken from female mosquitoes and observed under a high-resolution JEM 2-100 transmission electron microscope (JEOL) at an operating voltage of 80 kV. Raw TEM images were analyzed in Image J version 1.52e.^60^ Femur leg sections were taken from five random mosquitoes of each populations/strain. In total, 25 and 32 sections were measured for SEK and ASN, respectively. A Mann-Whitney *U* test (R-framework) was used to test for a significant difference in the thickness of the leg procuticle.

## 3. Results

### 3.1. RNA sequencing

Illumina sequencing generated ∼ 95-110 million strand-specific, paired-end reads per sample. Alignment of RNAseq reads against the *An*. *arabiensis* annotation resulted in an overall percent alignment rate of 89.2±0.7 (mean ± standard error of the mean, SE) across all samples (Table S2).

### 3.2. Principal Component Analysis (PCA)

A PCA using the 1000 most variable genes across all RNAseq samples revealed that 34.7 % of the total variation could be explained by PC1 while 32.8 % could be explained by PC2 (Figure 1). RNAseq replicates clustered by population/strain, either on PC1 (SEK) or both PC1 and PC2 (ASN, CHW, MOZ and TOL). RNAseq replicates of two resistant *An. arabiensis* populations, ASN and TOL, clustered together and away from those of the two susceptible strains (SEK and MOZ) while RNAseq replicates of the third resistant population, CHW, clustered between RNAseq replicates of ASN/TOL populations and RNAseq replicates of the susceptible SEK strain.

**Figure 1.**
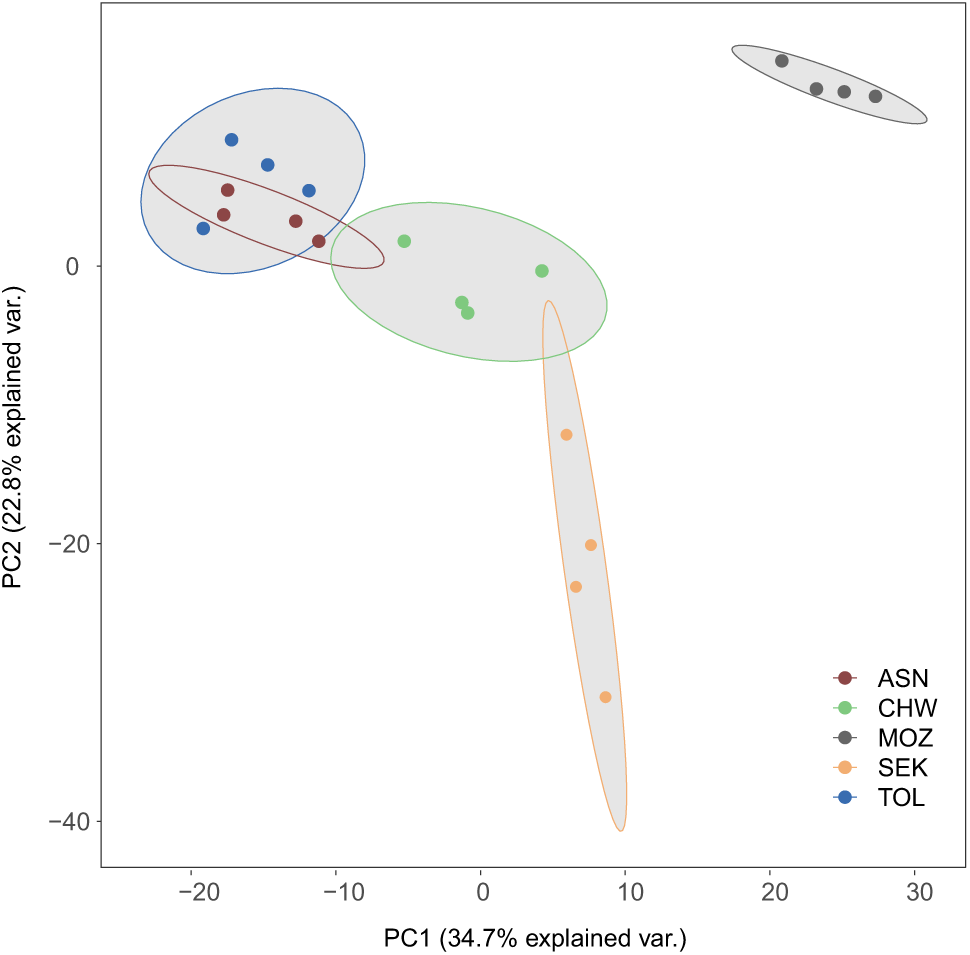
PCA analysis of gene expression among insecticide resistant and susceptible populations or strains. Three Ethiopian deltamethrin/DDT resistant populations (ASN, CHW and TOL) and two susceptible strains (SEK and MOZ) of *An. arabiensis* are as indicated.

### 3.3. Differential gene expression analysis

We used DESeq2 to perform a differential gene expression (DE) analysis ((foldchange (FC) ≥ 2 and a FDR < 0.05) between each resistant *An. arabiensis* population (ASN, CHW or TOL) and each of the susceptible *An. arabiensis* strains (SEK or MOZ). 496, 152 and 602 genes were overexpressed by two-fold or more, while 286, 109 and 197 *An. arabiensis* genes were underexpressed by twofold or more in ASN, CHW and TOL compared to the susceptible strain SEK, respectively. 936, 460 and 814 genes were overexpressed by twofold or more, while 798, 576 and 654 genes were underexpressed by twofold or more in ASN, CHW and TOL compared to the susceptible strain MOZ, respectively (Figure 2, Table S3). Not surprisingly, the total number of DEGs was lower for the DE analyses between one of the resistant populations (ASN, CHW and TOL) and a susceptible strain from the same country of origin (SEK, Ethiopia) compared to DE analyses using a susceptible strain from a different country of origin (MOZ, Mozambique). Inspecting the overlap of DEGs between the two DE analyses (against either SEK or MOZ) performed for each resistant population revealed that 303, 66 and 337 genes were overexpressed and 48, 14 and 29 genes were underexpressed in ASN, CHW or TOL compared to both SEK and MOZ, respectively. Furthermore, thirty-eight and four DEGs (hereafter named “core” DEGs) were over‐ and underexpressed, respectively, in each resistant population and for each comparison (Figure 2, Figure 3). The 38 overexpressed “core” DEGs coded for 14 uncharacterized proteins, 13 cuticle related proteins (either with an “insect cuticle protein” domain (IPR000618), a “chitin binding” domain (IPR002557) or defined as a “cuticle protein” by VectorBase), 2 nicotinic Acetylcholine Receptor subunits (nAChRs), Yellow-e, chitin synthase, GSTD3, a protein with a protein kinase domain (IPR011009), a nuclear-pore complex protein, a pyroglutamyl-peptidase, a thioester containing protein (tep1), a serine-type endopeptidase and a vitamin K-dependent protein C-like. The four underexpressed “core” DEGS coded for an uncharacterized protein, a protein (FBN8) with a fibrinogen domain (InterPro domain IPR002181), a G-protein coupled receptor (GPCR) and a dynein assembly factor. Of particular note, in line with the PCA in which the replicates of the resistant CHW population clustered most closely to those of the SEK strain, the CHW population had the lowest number of DEGs (against either SEK or MOZ) and almost all “core” DEGs (34/38) had a lower fold change in the CHW comparisons then in ASN or TOL comparisons (Figure 3). Finally, the fold changes of a selection of DEGs determined by DE analysis was shown to be consistent with those obtained by RT-qPCR (Figure 4, Figure S2).

**Figure 2.**
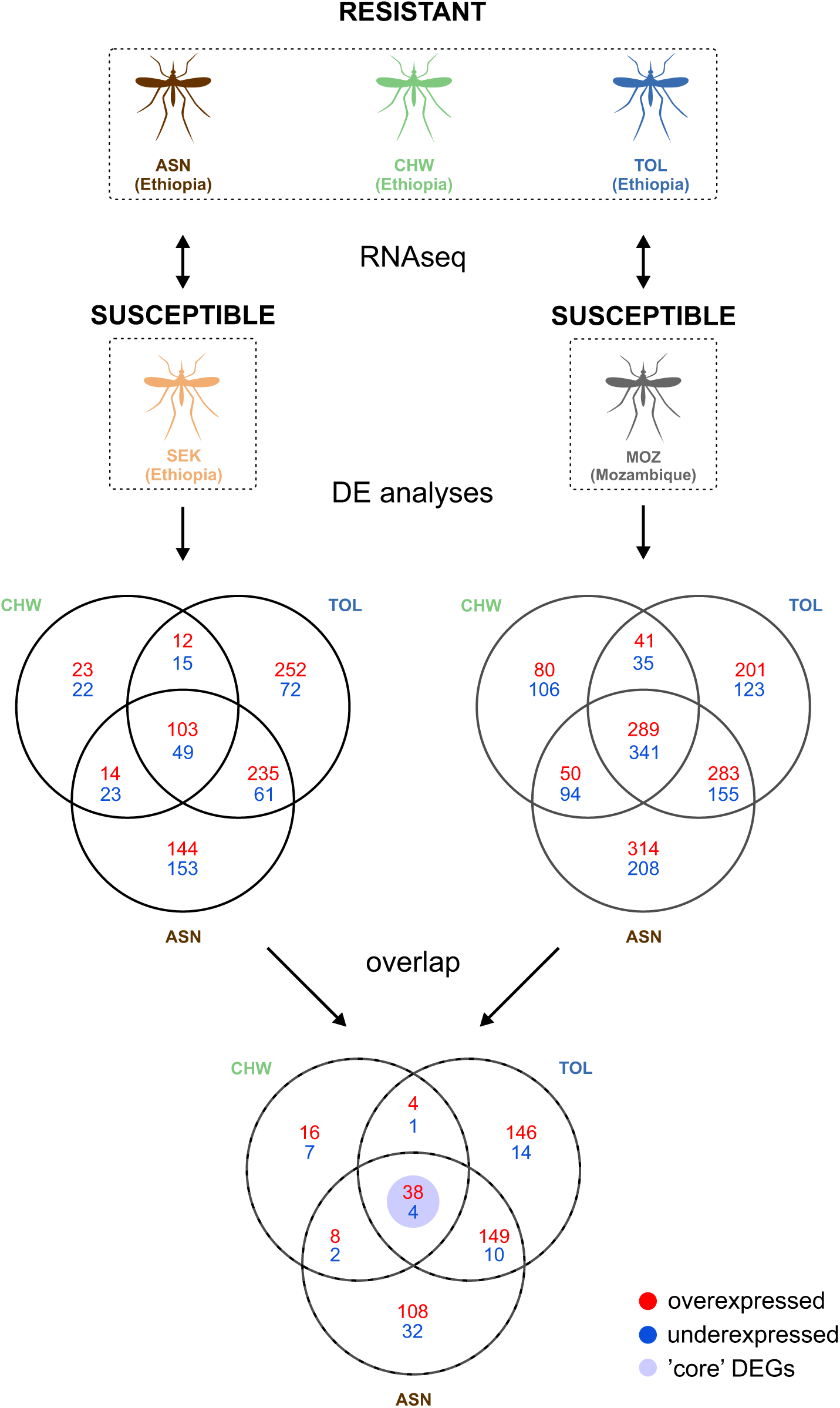
Experimental design and DEGs between three Ethiopian deltamethrin/DDT resistant populations and two susceptible strains of *An. arabiensis*. Differential gene expression was assessed between each resistant population (ASN, CHW or TOL) and each susceptible strain (SEK or MOZ) (FDR of 0.05, |log2 FC change| ≥ 1). Genes differentially expressed in each comparison of a deltamethrin/DDT resistant population against a suscepible strain are referred to as ‘core’ differentially expressed genes (‘core’ DEGs). For a list of all DEGs for each comparison, see Table S3.

### 3.4. GO enrichment analysis

A Gene Ontology (GO) enrichment was performed for each DE analysis using GOseq. A list of over‐ and underrepresented GOs for each comparison (six in total) can be found in Table S4. Those GO Molecular Function terms that were significantly overrepresented in both comparisons of a resistant population (against SEK or MOZ) are shown in Figure 5. For at least five out of 6 comparisons the GO-terms “structural component of the cuticula” (GO:0042302), “chitin-binding” (GO:0008061), “serine type endopeptidase” (GO:0004252) and “heme” (GO:0020037) were significantly overrepresented, while “oxidoreductase activity” (GO:0016705), “monooxygenase activity” (GO:0004497) and “iron ion binding” (GO:0005506) were overrepresented in at least one DE analysis of a resistant population. The GO enrichment results are also reflected in an expression heatmap of cuticle related genes, with clear expression pattern differences between the comparison of resistant populations against SEK and the comparison of MOZ against SEK (Figure S3).

### 3.5. Gene expression levels of P450s and GSTs in deltamethrin/DDT resistant *An. arabiensis* populations

All significantly overexpressed genes (FDR < 0.05, Table S3) were mined for members of detoxification gene families known to be involved in metabolic resistance against pyrethroids (P450s and GSTs, see Introduction). Only *gstd3* was significantly overexpressed in each comparison of a resistant *An. arabiensis* population against one of the susceptible strains (SEK or MOZ) (Figure 3, Table S3). Next, we investigated the expression level of genes encoding *Anopheles* P450s and GSTs known to metabolize pyrethroids (CYP6M2, CYP6P3, CYP6P4 and GSTE2^25, 26, 31, 36^). *Cyp6m2*, *cyp6p3* and *cyp6p4* were significantly overexpressed in all comparisons against MOZ (log_2_FC ranging from 2.2 to 3.6). *Cyp6p4* was significantly overexpressed in the comparison of each resistant strain against SEK, *cyp6p3* was significantly overexpressed in the comparison of CHW against SEK (log_2_FC of 1.0) while *cyp6m2* was not significantly overexpressed in any of the comparisons against SEK. *Gste2* was significantly overexpressed in CHW against MOZ and in the comparisons of ASN and TOL against either SEK or MOZ (Table S3). A similar trend could be observed for the expression values obtained by RT-qPCR, with fold changes of *cyp6m2*, *cyp6p3, cyp6p4* and *gste2* being higher in the comparisons against MOZ compared to comparisons against SEK (Figure 4). Furthermore, we also evaluated the expression of *cyp4g16*, a gene encoding a P450 catalyzing epicuticular hydrocarbon biosynthesis. This gene was significantly overexpressed in all comparisons against MOZ and SEK. RT-qPCR data confirmed *cyp4g16* overexpression in the case of ASN or TOL versus SEK. Finally, both RNAseq data and RT-qPCR data showed that *cyp4c28*, a P450 gene previously shown to be overexpressed in resistant *Anopheles* sp. ^43, 61^, was significantly overexpressed in all comparisons against SEK, but not against MOZ (Figure 4)

**Figure 3.**
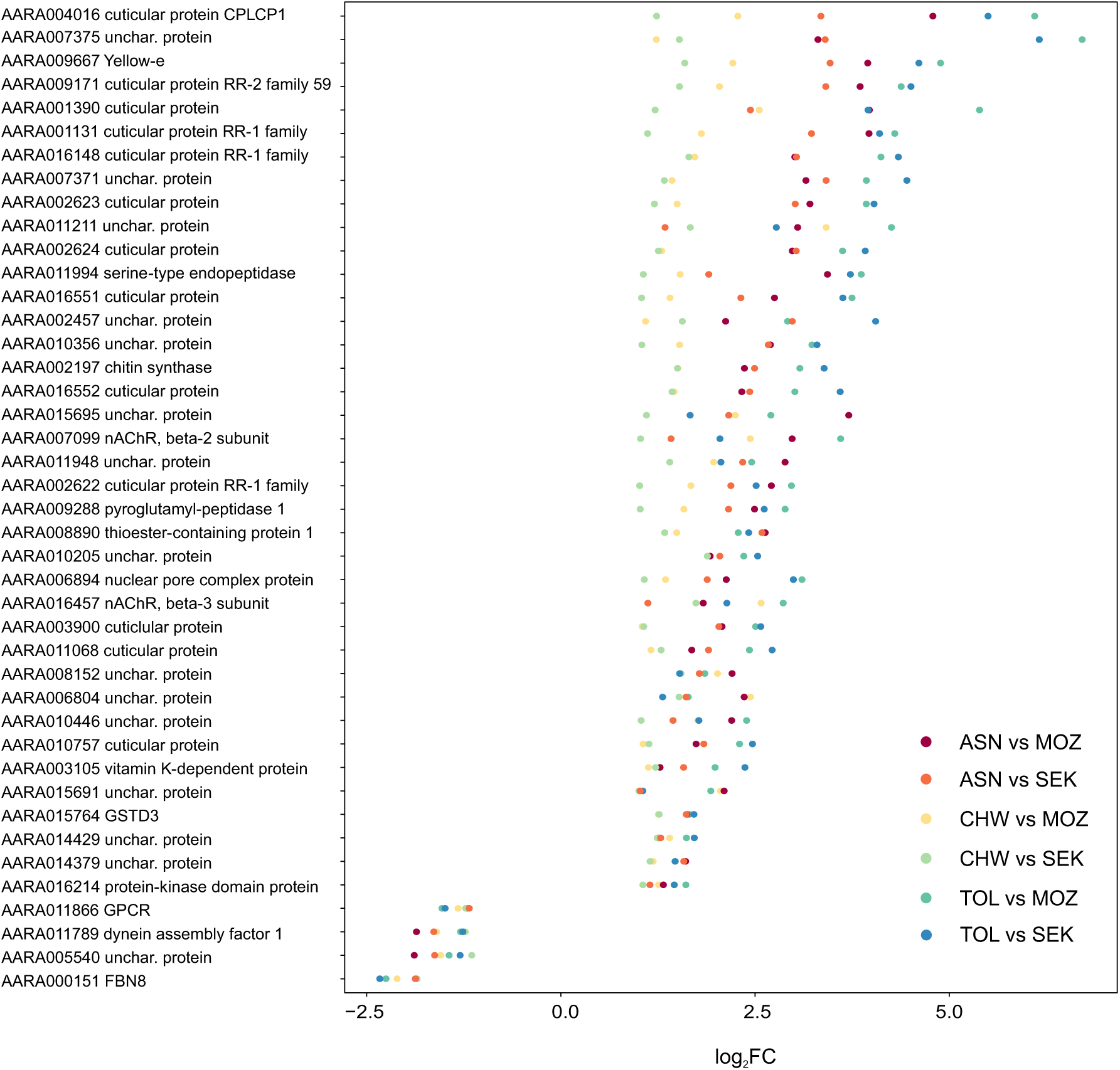
Identity of *An. arabiensis* ‘core’ DEGs and their fold change between Ethiopian deltamethrin/DDT resistant populations (ASN, CHW or TOL) and two susceptible strains (SEK or MOZ).

### 3.6. Detection of mutations involved in insecticide resistance

The RNAseq reads of all resistant populations and susceptible strains were mined for mutations involved in resistance against either DDT, pyrethroids, cyclodienes and organophosphates (Table S5). None of the known *gste2* and *AChE1* resistance mutations could be identified in the RNAseq reads of the populations/strains of this study. The L1014F mutation in the *vgsc* (A2532T, codon change of TTA to TTT, in the coding sequence of AARA016386-RA) was identified in all resistant populations (ASN, CHW and TOL) and the susceptible SEK strain. Finally, the A301S mutation in *Rdl* (G886T, codon change of GCA to TCA, in the coding sequence of AARA016354-RA) was identified in two resistant populations (CHW and ASN) and in the susceptible SEK strain (Table S5).

### 3.7. Procuticle thickness in mosquito legs

The leg procuticle thicknesses of the deltamethrin/DDT resistant population (ASN) and the susceptible strain (SEK) were 2.40±0.12 μm and 2.41±0.06 μm (mean ± SE), respectively, and were not significantly different (p > 0.05) (Figure 6).

## 4. Discussion

*An. arabiensis* is one of the dominant vector species of malaria in sub-saharan Africa including Ethiopia^62^, where resistance of *An. arabiensis* against pyrethroids and DDT is widespread.^10^ In many Ethiopian *An. arabiensis* populations an association between the *kdr* mutation in the VGSC and resistance to pyrethroids and DDT resistance has been observed. ^7, 9, 10, 12^ However, a number of studies have also pointed to increased detoxification as important in resistant populations, as the resistance phenotype is strong, and because *kdr* mutations were not fixed at the population level.^10, 12^ Metabolic resistance has been observed in other pyrethroid and DDT resistant *An. arabiensis* populations from East-Africa, but the putative involvement of genes encoding detoxification enzymes associated with metabolic resistance was only investigated for populations from Tanzania and Sudan using a whole-genome microarray.^32, 42, 43^ In this study, we expand our previous work on resistance monitoring of Ethiopian *An. arabiensis* populations^12^ and used Illumina sequencing to quantify gene expression levels in deltamethrin and DDT resistant *An. arabiensis* populations from three different sites in Ethiopia ‐ Asendabo (ASN), Chewaka (CHW) and Tolay (TOL) and in two deltamethrin and DDT susceptible laboratory strains, MOZ and SEK (Figure 2).

First, we mined Illumina RNAseq data to assess the prevalence of mutations previously associated with insecticide resistance in *Anopheles* species. We also estimated the frequency of these mutations, but as both gene expression and allele-specific gene expression can significantly influence the accuracy of allele frequency estimation^41, 63^ we did not integrate these results into the discussion section of this study. None of the populations harbored mutations in the *gste2* gene nor in the *AChE1* gene, but in both ASN, CHW and SEK an A301S mutation in the *Rdl* gene, associated with resistance against cyclodienes, could be identified (Table S5, see also Introduction). The presence of the A301S mutation most likely reflects the long historical use of cyclodienes in malaria vector control.^64^ In addition, although Ethiopia banned cyclodienes in 2004^65^, cyclodienes such as endosulfan and chlordane are still detectable in environmental samples from some regions of Ethiopia.^66, 67^ In line with Alemayehu *et al.* 2017, all three resistant populations (ASN, CHW and TOL) harbor the *kdr* mutation L1014F while the N1575Y mutation was absent.^12^ We also identified the L1014F mutation in the deltamethrin and DDT susceptible SEK strain (Table S5).

Because target site mutations are unlikely to fully explain high-level resistance in Ethiopian populations (^10, 12^ and see above), we performed a differential gene expression analysis between each deltamethrin and DDT resistant *An. arabiensis* population (ASN, CHW or TOL) and both susceptible strains (SEK or MOZ) (Figure 2). The differential expression was more pronounced, both in number of DEGs and in magnitude of differential expression, for the comparison of the Ethiopian resistant populations against MOZ (Figure 2, Table S3). This might reflect genetic (and expression) variation by distance, as the MOZ strain originated from Mozambique, while SEK is a strain from Ethiopian. Consistent with a role in resistance, we found that members of detoxification gene families known to be involved in metabolic resistance of anopheline mosquitoes against pyrethroids and/or DDT varied in expression in our study. According to the RNAseq and/or RT-qPCR data, *cyp6p4* was significantly overexpressed in the resistant strains (ASN, CHW or TOL) compared to any of the susceptible strains (MOZ or SEK) (Figure 4, Table S3). Recently, it has been shown that CYP6P4 is the major P450 responsible for pyrethroid resistance in a *kdr*-free population of *An. arabiensis* from Chad. However, although it was shown that this P450 could metabolize several Type I and Type II pyrethroids, it could only bind to deltamethrin and not metabolize this compound. ^31^ Thus, the overexpression of *cyp6p4* in ASN, CHW and TOL might be related to resistance against pyrethroids other than deltamethrin. It remains to be tested whether resistance to such pyrethroids (e.g., permethrin and lambda-cyhalothrin) is present in these Ethiopian populations.^12^ Only in CHW, *cyp6p3* and *cyp6m2* were significantly overexpressed as compared to both susceptible strains (Figure 4, Table S3). Previously, *An. gambiae* CYP6P3 and CYP6M2 were shown to metabolize deltamethrin and/or DDT, and hence the overexpression of their orthologue in CHW might contribute to metabolic resistance against these insecticides. ^25, 26, 29^ In 2014, Riveron *et al.* showed that *An. funestus* GSTE2 was able to metabolize DDT. For both ASN and TOL, *gste2* was significantly overexpressed compared to both susceptible strains, suggesting this enzyme might also play a role in metabolic DDT resistance in Ethiopian *An. arabiensis* populations. Last, *gstd3* was significantly overexpressed for each comparison of a resistant population against a susceptible strain (Figure 3, Table S3). *Gstd3* overexpression has been reported for several pyrethroid and DDT resistant *Anopheles* populations^35, 68, 69^, but at present the role of delta class GSTs is thought to be minor compared to those of the epsilon class (e.g., GSTE2, see above) and functional validation of the interaction between GSTD3 and DDT/deltamethrin is needed to understand the contribution of this GST towards DDT/deltamethrin resistance.^68^

**Figure 4.**
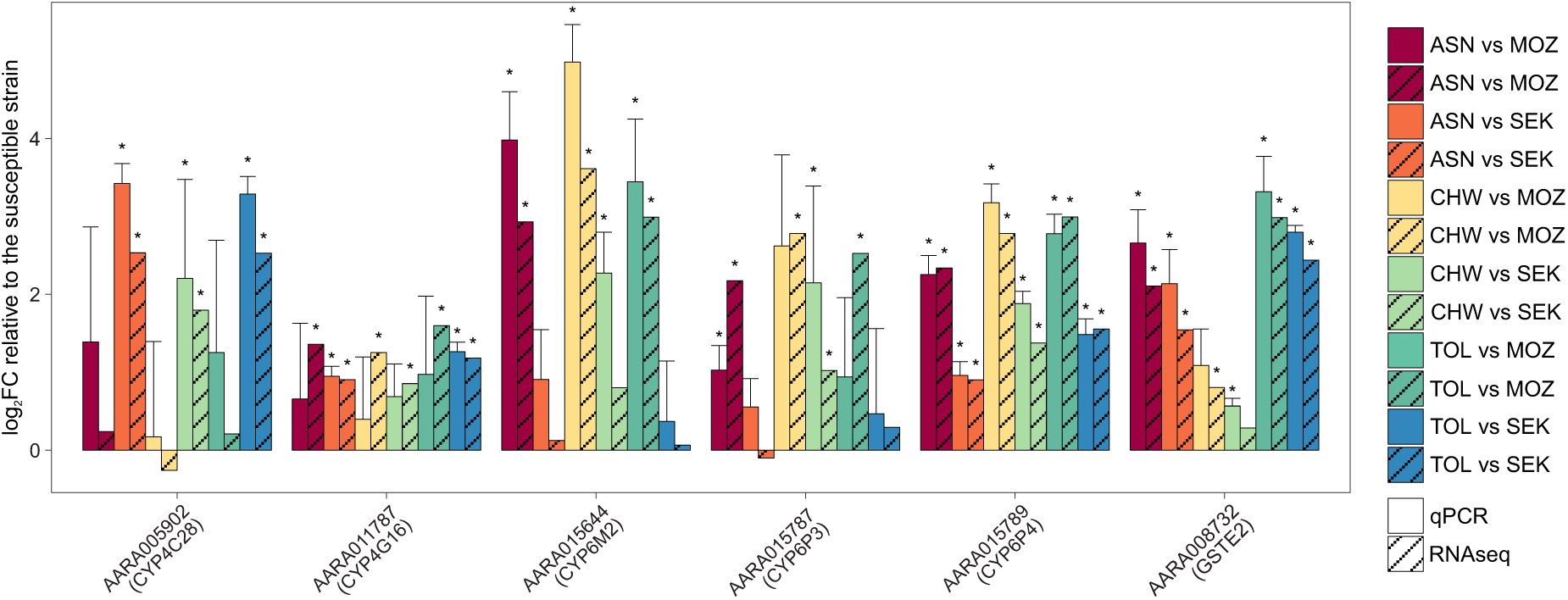
Expression levels of GST and P450 genes in Ethiopian deltamethrin/DDT resistant populations (ASN, CHW or TOL) compared to two susceptible strains (SEK or MOZ) of *An. arabiensis*. An asterisk indicates whether a P450 or GST gene is significantly overexpressed, either based on RNAseq (FDR of 0.05, Table S3) or RT-qPCR data (student’s unpaired t-test *p*-value < 0.05).

Including *gstd3*, 41 genes belonged to the “core DEGs” set that were differentially expressed in each resistant population and for each comparison (against SEK or MOZ). Thirteen (32%) of these 41 genes encode cuticular proteins that are overexpressed, while others encode chitin synthase, yellow-e protein, serine-type endopeptidase, uncharacterized proteins and two nicotinic acetyl-choline receptors (AChRs) beta subunits (Figure 3). It has been shown that pyrethroids exert (secondary) non-specific inhibitory effects on nicotinic AChRs^70^ and as such their upregulation in the resistant *An. arabiensis* strains might be a way to compensate for non-specific nAChR inhibition. To complement our set of “core DEGs”, we also performed a GO analysis for each DE comparison (Figure 5). In agreement with the expression analysis of the major detoxification genes involved in deltamethrin/DDT resistance (see above), GO-terms related to P450 activity were significantly enriched in at least one of the different DEG sets. In addition, also in line with our set of “core DEGs”, three GO-terms related to changes in the cuticula were significantly enriched in nearly every DEG set (Figure 5). This is also reflected in a heatmap of expression changes of cuticle related genes in deltamethrin/DDT resistant populations ASN, CHW and TOL, as shown in Figure S3. Higher expression of cuticular genes has previously been reported for pyrethroid resistant mosquito populations^32, 71–75^ and in some cases was associated with a thicker cuticula.^73, 74^ Further, some of these *Anopheles* cuticular genes were also shown to be expressed in the limbs, the most frequent site of contact with insecticides.^76^ Apart from genes encoding cuticular proteins, *cyp4g16*, which encodes a P450 that catalyzes epicuticular hydrocarbon biosynthesis, has also been reported to be frequently overexpressed in insecticide resistant *Anopheles* mosquitoes, including *An. arabiensis*^32–34, 42, 73, 77^. This has led to the suggestion that CYP4G16 plays a role in insecticide resistance via enrichment of the cuticular hydrocarbon (CHC) content.^33^

**Figure 5.**
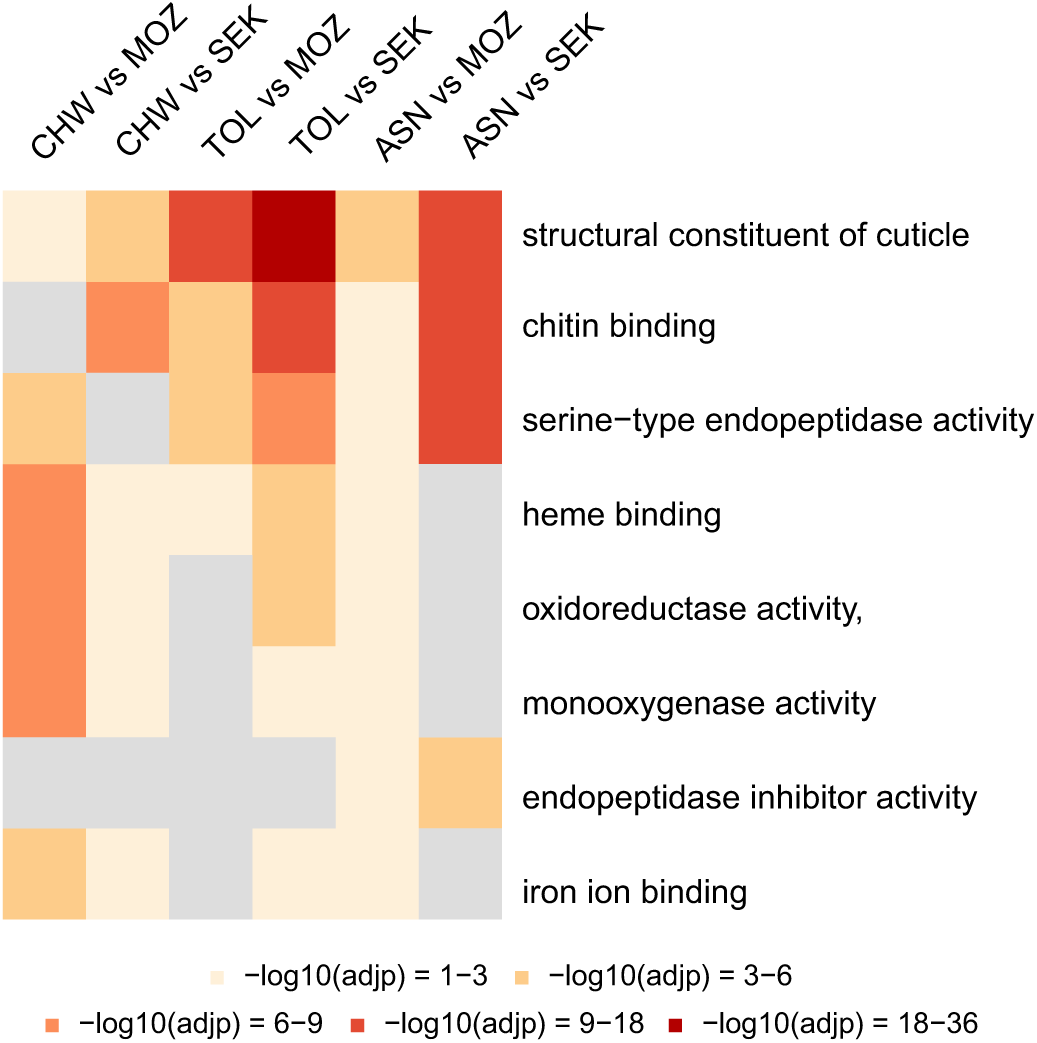
GO enrichment analysis of DEGs in three Ethiopian deltamethrin/DDT resistant populations (ASN, CHW or TOL) compared to two susceptible strains (SEK or MOZ) of *An. arabiensis*. Heatmap showing the FDR of GO categories among DEGs of each comparison of a resistant population against a susceptible strain. A grey colored cell indicates that the GO category was not significantly enriched (FDR ≥ 0.05) for a given comparison. Only GO Molecular Function terms that were significantly overrepresented in both comparisons of a resistant population against SEK and MOZ are shown.

According to our differential expression analysis (RNAseq and/or RT-qPCR data, see Figure 4 and Table S3), *cyp4g16* was overexpressed as compared to both susceptible strains in the ASN and TOL populations. Both mechanisms (a greater cuticle thickness by cuticle protein overexpression or CHC enrichment of the epicuticle) might reduce the penetration rate of insecticide and may enhance resistance by increasing the time available for metabolic processes to inactivate the insecticide before it causes target-site inhibition. We therefore measured the thickness of the procuticle (comprising an exo-, meso‐ and endocuticle)^78^ of a representative resistant population (ASN) and a susceptible strain (SEK). In contrast to Yahouédo et al. (2017), who found that the procuticle of a resistant *An. gambiae* strain was thicker than that of a susceptible strain^73^, we did not detect a statistical difference between the average leg procuticle thickness of the resistant population and susceptible strain of this study (Figure 6). However, in Balabanidou *et al.* 2016 the epicuticle, layered on top of the procuticle, was the main contributor to differences in cuticle thickness between pyrethroid resistant and susceptible populations.^33^ Unfortunately, we were not able to measure epicuticle thickness in this study as the epicuticle was not preserved in the majority (> 95%) of the *An. arabiensis* leg sections. Alternatively, it could be that the epicuticle of the resistant Ethiopian *An. arabiensis* populations has a higher CHC content compared to those of the susceptible strains, and hence determining CHC levels in both resistant populations and susceptible strains merits further investigation. On the other hand, in contrast to altered thickness or CHC levels, it could be that a change in composition of the cuticle is associated with deltamethrin/DDT resistance, as reviewed by Balabanidou *et al.*^34^ For example, a gene encoding a laccase with a key-role in sclerotization was overexpressed in a resistant *Culex* population.^79^ Strikingly, in our study, the gene *yellow-e* was overexpressed in each comparison of a resistant population to each of the susceptible strains (Figure 3). In *Tribolium castaneum*, YELLOW-E was shown to have an important role in cuticle pigmentation/tanning^80^ and, hence, its overexpression in *An. arabiensis* populations might lead to an altered cuticle, possibly reducing the penetration rate of deltamethrin or DDT. Future work should study the role of cuticle composition as a potential resistance factor in Ethiopian populations of *An. arabiensis*.

**Figure 6.**
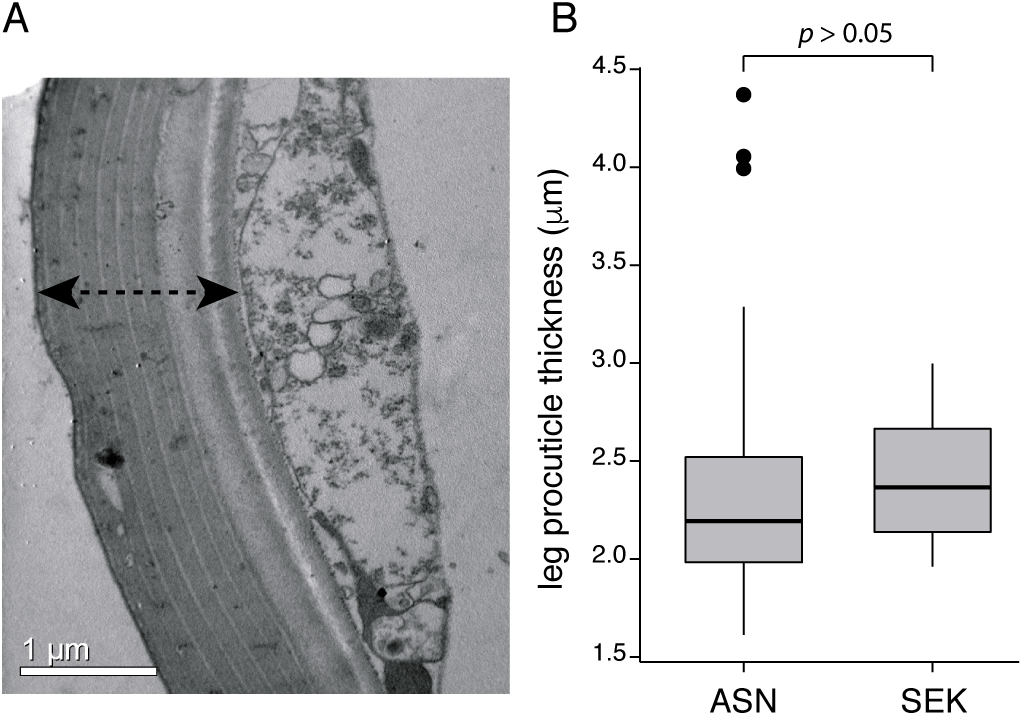
Leg procuticle thickness does not differ between a deltamethrin/DDT resistant population and a susceptible strain of *An. arabiensis*. A: Representative image of a cross section of the femur leg segment (SEK strain). Only the procuticle (indicated by a double headed arrow) was measured as the epicuticle was not preserved during preparation of sections. B: Box plot showing the distribution of leg procuticle thickness measurements of the deltamethrin/DDT resistant population (ASN) and the deltamethrin/DDT susceptible strain (SEK). Outliers are represented as black circular dots. Distributions were compared using a Mann-Whitney *U* test.

## Author Contributions

ES, TVL and LD conceived and designed study. ES and VB performed experiments. WD, ES, SS and AB analyzed data. WD and ES wrote the manuscript, with input from JV, RMC, DY, LD and TVL. All authors read and approved the final manuscript.

## Acknowledgements

The authors are grateful to Flemish Interuniversity Council (VLIR-UOS) for the financial support of this study. We thank Robert Greenhalgh for assistance in managing Illumina read data and would like to acknowledge the assistants of the Tropical and Infectious Diseases Research Center (TIDRC) for their technical support during field and preliminary lab work. WD is a postdoctoral fellow of the Research Foundation-Flanders (FWO). This project was funded by the Research Foundation Flanders (FWO, Belgium) (grant G009312N to TVL and grant G053815N to TVL and WD) and by the European Union Horizon 2020 Framework Program (688207-DMC-MALVEC) to JV and DY. VB was supported by a Scholarship for Strengthening Post-Doctoral Research from The Greek State Scholarships Foundation (IKY) within the framework of the Operational Programme “Human Resources Development Program, Education and Life-Long Learning”. Research reported in this publication utilized the High-Throughput Genomics and Bioinformatic Analysis Shared Resource at Huntsman Cancer Institute at the University of Utah and was supported by the National Cancer Institute of the National Institutes of Health under Award Number P30CA042014. The content is solely the responsibility of the authors and does not necessarily represent the official views of the funding agencies.

## Supporting Information

**Table S1.** List of selected candidate genes for RT-qPCR validation and the used qPCR primer sequences.

**Table S2.** Read statistics for RNAseq samples of Ethiopian deltamethrin/DDT resistant populations (ASN, CHW and TOL) and two susceptible strains (SEK and MOZ) of *An. arabiensis*.

**Table S3.** Differentially expressed genes between Ethiopian deltamethrin/DDT resistant populations (ASN, CHW and TOL) and two susceptible strains (SEK and MOZ) of *An. arabiensis*.

**Table S4.** GO enrichment analysis of differentially expressed genes between Ethiopian deltamethrin/DDT resistant populations (ASN, CHW and TOL) and two susceptible strains (SEK and MOZ) of *An. arabiensis*.

**Table S5.** Ratio of resistance mutations in Ethiopian deltamethrin/DDT resistant populations (CHW, ASN and TOL) and two susceptible strains (SEK and MOZ) of *An. arabiensis.*

**Figure S1.** Map of Ethiopia showing the collection sites of the three deltamethrin/DDT resistant *An. arabiensis* populations.

**Figure S2.** RT-qPCR validation of differentially expressed genes between Ethiopian deltamethrin/DDT resistant populations (ASN, CHW and TOL) and two susceptible strains (SEK and MOZ). A tilde (∼) indicates cuticle related genes. For a description of each gene see Table S1.

**Figure S3.** Expression heatmap of cuticle related genes of *An. arabiensis*

Cuticle related genes were defined as those genes coding for proteins with one of the following InterPro domains: IPR000618, IPR031311, IPR31874, IPR002557, IPR22727, IPR004302 or IPR004835. The log_2_ transformed gene fold changes of the Ethiopian deltamethrin/DDT resistant populations ASN, CHW, TOL and the susceptible strain MOZ from Mozambique are relative to the susceptible SEK strain from Ethiopia. Genes without expression values in all four comparisons were excluded from the heatmap. *Anopheles arabiensis* gene IDs are shown on the right.

**File S1.** Gene Transfer Format (GTF) used for mapping and counting of *An. arabiensis* RNAseq reads.

